# Humanized Kcnv2 E151X Mouse Captures Hallmarks of KCNV2-Associated Retinal Dystrophy

**DOI:** 10.64898/2026.01.05.697593

**Authors:** Nermina Xhaferri, Sumit Biswas, Benjamin Davies, Moritz Lindner

**Affiliations:** Institute of Physiology and Pathophysiology, Department of Neurophysiology, Philipps-University Marburg, Deutschhausstrasse 2, 35037 Marburg, Germany; Francis Crick Institute, London, Great Britain; Department of Ophthalmology, Marburg, University Hospitals of Giessen and Marburg

**Keywords:** KCNV2, KCNV2- associated retinopathy, Inherited retinal dystrophy, photoreceptors, electroretinography

## Abstract

**Background:** KCNV2-associated retinopathy is a rare inherited retinal dystrophy caused by variants in the KCNV2 gene, leading to disrupted photoreceptor function and slowly progressive vision loss. Patients have characteristic electroretinography abnormalities, including reduced cone responses, delayed and reduced rod responses to low light flashes and paradoxally large rod-driven responses to bright flashes of light. To model this condition, we generated the Kcnv2 E151X mouse line and assessed its structural and functional retinal features.

**Methods:** We have employed CRISPR/Cas 9 gene editing technology to generate a mouse line with an early stop mutation in position E151- orthologous to the commonly encountered E143X mutation in humans - and performed a combination of Iimmunohistochemistry and Western Blot to confirm the absence of the full-length KCNV2-encoded protein, K_v_8.2. To assess how closely it models the human disease, we have characterised the KCNV2 mutant mouse line at histological and functional levels employing immunohistochemistry and electroretinography, respectiveley.

**Results:** Kcnv2 mutant mice showed markedly reduced photopic responses and reproduced the supernormal rod phenotype as described in affected individuals. In the morphological context, mutant retinas demonstrated strong and early glial fibrillary acidic protein upregulation together with reduced counts of cone arrestin positive cells as well as photoreceptors in general. Power calculations based on the data obtained herein suggest therapeutic trials are feasible with small sample sizes.

**Conclusions:** The Kcnv2 mutant mouse line replicates key functional and structural hallmarks of KCNV2– associated retinopathy. This model provides a relevant platform for mechanistic studies and preclinical evaluation of gene based or pharmacological therapies targeting cone and rod photoreceptor dysfunction.

## Introduction

KCNV2- associated retinopathy, previously known as cone dystrophy with supernormal rod response (CDSRR), is an inherited retinal disorder characterized by a slowly, progressive deterioration of visual function. It was first described in 1983 by Gouras et al. [1] and subsequently linked to mutations in the KCNV2 gene [2, 3]. This encodes for a voltage-gated potassium channel subunit, K_v_8.2, which is expressed in rod and cone photoreceptors. While KCNV2- associated retinopathy is overall rare, it is one of the commonest causes of autosomal-recessive cone dystrophies [4] and the slow rate of disease progression leaves ample room for therapeutic intervention. [5]

A recent study on mouse and primate photoreceptors [6] has shown that K_v_8.2 interacts with K_v_2.1 in rod inner segments and with both K_v_2.1 and K_v_2.2 in cone inner segments, giving rise to I_K,x_, the conductance that physiologically counterbalances the constant influx of positive charge through CNG channels (“dark-current”) in resting photoreceptors. K_v_2.1 homomers and K_v_8.2/K_v_2.1 heteromers differ in their voltage-dependence of activation and inactivation. Heteromeric K_v_8.2/K_v_2.1 channels typically activate and inactivate at more hyperpolarized potentials than K_v_2.1 homomers. Moreover, the total conductance of K_v_8.2/K_v_2.1 channels is lower than that of K_v_2.1 homomers and the activation time is shortened.[2, 7, 8] The change in electrophysiological properties in the absence of K_v_8.2 translates to distinct features in the electroretinograms of the patients including the diagnostic supranormal scotopic b-wave response to bright flash. Other typical electroretinographic observations include delayed and decreased scotopic responses to low-flash intensities, and markedly reduced to absent photopic responses. Another characteristic finding is the broadening of the a-wave trough following scotopic bright flash stimulation. [3, 9–12]

Different reports collectively highlight the diversity of pathogenic Kcnv2 mutations including a variety of nonsense (E73X, Q76X, E143X, E148X, K260X, Q287X), missense (E184K, E184V, G461R), frameshift (K120fsX371) and deletions (D339_V341del, gross deletions). [3, 10, 13, 14] Thereof, E143X is the commonest single pathogenic mutation [15] leading to an early truncation of K_v_8.2 inside the tetramerization domain [16].

Patients with KCNV2 retinopathy typically present in the first two decades of their life with decreased visual acuity, color vision abnormalities, photophobia and night blindness. As shown by fundoscopy and clinical imaging, structural alterations in cones are observed early in the course of the disease while the rod-dominated peripheral retina appears generally normal. [3, 17]

Despite the growing insights into the genetic underpinnings of KCNV2 retinopathy, the precise molecular mechanisms by which KCNV2 mutations lead to retinal dysfunction and ultimately cone death remain poorly understood. Critical questions still remain unanswered: How do KCNV2 mutations affect I_k,x_? What are the molecular pathways driving retinal cell death and why are cones predominantly affected? Could a medical or gene-therapeutic intervention aiming to restore physiological I_K,x_ only alleviate symptoms or also prevent cone loss?

In this study, we have generated and characterized a novel mouse model carrying the E151X mutation, orthologous to E143X in humans, not only to address what happens at the basis of the disease with a model that better resembles human disease cases than previous models, but also to evaluate its potential utility as a platform for developing novel therapeutic approaches. Via a combination of immunohistochemistry, Western Blot and electroretinogram experiments we have shown that these mice closely mimic the phenotype of KCNV2– associated retinopathy patients and provide a good model to elucidate the pathophysiological mechanisms of this blinding disease and pave the way for gene therapy strategies.

## Materials & Methods

### Animal procedures

All procedures were performed with the approval of the Giessen Regional Council Animal Health Authority (File No: G19/2023) and the UK Home Office (PAA2AAE49), respectively, and in accordance with the ARVO Statement for the Use of Animals in Ophthalmic and Vision Research. Wild-type C57BL/6J mice and Kcnv2 mice (Kcnv2 WT, Kcnv2^E151X/E151X^, Kcnv2^WT/E151X^) aged from 1 month to 1 years old, both males and females, were used in this study. Wild-type C57BL/6J mice were obtained from Charles River Laboratories (Sulzfeld, Germany) and Kcnv2 mice were an in-house breeding colony. Animals were housed in a temperature-controlled environment with a 12-hour light/dark cycle with no restriction on food and water. Mice were back-crossed for at least four generations before being used in experiments.

### Generation of Kcnv2E151X mice

Generation of the mouse line carrying the E143X mutation in the *Kcnv2* (NM_183179.1) gene (*Kcnv2^em1Mlin^, Kcnv2^E151X^ hereafter)* following established procedures [18] utilizing CRISPR/Cas9 gene editing technology. In brief, a target site (5’- GTGCCTCGTAGTCATCACAC-3’) for the nuclease on exon 1 (ENSMUSE00000395161) of the *Kcnv2* gene, overlapping the E151 residue, was identified using the CRISPOR design algorithm (http://crispor.tefor.net/). A single guide RNA (sgRNA) was synthesized by Synthego (Redwood City, CA, USA) and assembled into a ribonucleoprotein complex with Cas9 nuclease. A 139 nt single-stranded oligodeoxynucleotide (ssODN) designed to introduce the E151X mutation (5’-AAACGCGCCTGGGTCGCCTGGCCACCTCCACCACTCGCAGAGGCCAGCTGGGtCTGTGTGATGACTACtAGGCACAGACAGACGAGTACTTCTTTGACCGTGACCCAGCGGTCTTCCAGCTCATCTACAACTTCTACAC-3’) was used as the template for homology-directed repair. The CRISPR/Cas9 RNP and ssODN were introduced into fertilized C57BL/6J zygotes via electroporation as previously described [18] and cultured overnight. The resulting two-cell stage embryos were surgically implanted into foster females. From the offspring, two founder mice which had successfully incorporated the E151X mutation were identified and subsequently bred with wild-type C57BL/6J mice to produce heterozygous F1 mice. The transmission of the E151X mutation was verified through Sanger sequencing.

### Genotyping

Mouse genomic DNA was extracted from ear notches or tail tips by incubation in 50 mM NaOH, for 1 – 1.5 hours at 95° C. Subsequently a 477 bp region within exon 1 of *Kcnv2*, encompassing the target codon, was amplified via PCR (DreamTaq PCR Master Mix, Thermo Fisher, Waltham, MA, USA), utilizing primers Kcnv2-F (5’- CTC ACA AGC CTC CTG GAT GG-3’ and Kcnv2-R (5’- TCC ATA GAA GCG CAT GTC CC-3’). PCR products were digested with BfaI (New England Biolabs, Cat. No. R0568S, Ipswich, MA, USA) yielding two fragments of 294 and 183 bp for the E151X-allele and one 477 bp fragment for the wild-type allele.

### Immunohistochemistry

For immunohistochemistry experiments tissue was collected at different timepoints spanning from 4 to 52 weeks of age. Following dark adaptation for at least 20 minutes, mice were anesthetized using 5% isoflurane in 0.2 L/min oxygen and then sacrificed by decapitation. Two different tissue fixation strategies were used. The long fixation protocol has been described previously [19]. In brief, eyes were carefully removed and fixed in 4% paraformaldehyde (PFA, ThermoFischer Scientific Cat. No. 28906 Waltham, MA, USA) in phosphate-buffered saline (PBS) for 24 hours and transferred to a 30% sucrose solution. The eyes were then embedded in an optimal cutting temperature (OCT) compound (Sakura, Finetek, Torrance, CA, USA) and cryopreserved using either dry ice or isopropanol at -40°C. The short fixation protocol involved removing the eyes and carefully extracting the retinas in ice-cold PBS. The extracted retinas were fixed in 4% PFA in PBS for 30 minutes and then placed in a 30% sucrose solution until fully drowned. Subsequently, the retinas were transferred to a mixture of 30% sucrose and OCT in a 3:1 ratio, before being embedded in undiluted OCT compound and cryopreserved as above. For both protocols, the OCT-embedded retinal sections were cut at a thickness of 20-25 μm, using a Leica CM1850 cryostat and stored at -20°C until further use.

Immunofluorescence staining was performed as previously described [20] using the antibodies and dilutions shown in table 1. Nuclei were counterstained with DAPI for 10 minutes. Finally, sections were mounted using Prolong Gold antifade mounting medium and coverslipped for imaging.

**Table 1.** List of primary and secondary antibodies used in this study. *Full aminoacid sequence of the immunizing peptide: MLKQSNERRWSLSYKPWSTPETEDVPNTGSNQHRRSICSLGARTGSQASIAPQWTEGNYNYYIEEDEDCGEEGEWKDDLAEENQKAECLTSLLDGHNDTPAQMSTLKVNVGGHSYLLECCELANYPKTRLGRLATSTTRRGQLGLCDDYEAQTDEYFFDRDP

| Target | Host | Source | Dilution |
| --- | --- | --- | --- |
| *Primary* |  |  |  |
| Kcnv2* | Mouse | Antibodies Incorporated Cat# 75-435, RRID: AB_2651161 | 1:400 |
| Kcnv2 | Rabbit | Sigma-Aldrich Cat# HPA031131, RRID: AB_10609996 | 1:400 |
| GFAP | Rabbit | Abcam Cat# ab7260, RRID: AB_305808 | 1:1000 |
| Cone Arrestin | Rabbit | Millipore Cat# AB15282, RRID: AB_1163387 | 1:500 |
| *Secondary* |  |  |  |
| Mouse | Donkey | Invitrogen Alexa Fluor 488 Cat#A-21202 RRID: AB_141607 | 1:1000 |
| Rabbit | Donkey | Invitrogen Alexa Fluor 488 Cat#A-21206 RRID: AB_2535792 | 1:1000 |
| Mouse | Donkey | Invitrogen Alexa Fluor 568 Cat#A-10037 RRID: AB_11180865 | 1:1000 |

### Imaging and Data analysis

Stained sections were imaged using a Zeiss LSM 710 confocal microscope (Zeiss, Oberkochen, Germany). Laser power, photomultiplier and digital gain, and pinhole size were kept constant for all samples belonging to the same experimental group, to ensure uniformity and comparability of acquired images. The obtained images were further processed and analyzed using ImageJ software (National Institute of Health, Bethesda, MD)[21]. This included global adjustment of brightness and contrast to optimize visualization. GFAP-positive fibrils were measured in ImageJ with the Segment tool, drawing along the line of each continuous GFAP-labeled segment from its visible origin at the ganglion cells level to its terminal tip. For outer nuclear layer thickness, we manually counted the number of contiguous rows of photoreceptor nuclei per field of view.

### Protein biochemistry and western blots

For protein biochemistry experiments, mice were sacrificed as described above. The retinas of Kcnv2 WT n=3), Kcnv2^WT/E151X^ (n=3), Kcnv2^E151X/E151X^ (n=3) mice were dissected in ice-cold PBS and transferred into Precellys lysing tubes (Bertin Technologies, Montigny-le-Bretonneux, France) containing 50 µl lysis buffer (1% triton X-100, 300mM NaCl, 0.4% sodium deoxycholate) and 0.5 µl protease inhibitor cocktail (Roche, Basel, Switzerland). The tissue was homogenized using the Minilys system and incubated with constant agitation for 30 minutes at 4 °C and then centrifuged. The supernatants were transferred to fresh tubes and the total protein concentrations were determined using the BCA method (Pierce BCA Protein Assay Kit, Thermo Fisher Scientific). Absorbance was measured at 562 nm using a NanoDrop OneC spectrophotometer. Each measurement was performed in duplicate. Protein concentrations were calculated based on standard curves. Equal amounts of protein (25 µg per sample) were mixed with sample loading buffer and incubated at room temperature for 30–60 minutes prior to electrophoresis. Samples, along with a molecular weight marker (Color Prestained Protein Standard, Broad Range, 10–250 kDa; New England Biolabs, Ipswich, MA, USA), were separated by sodium dodecyl sulfate–polyacrylamide gel electrophoresis (SDS-PAGE) on 10% gels. Proteins were transferred to nitrocellulose membranes using a wet transfer system. Membranes were blocked in 5% blotting-grade milk prepared in PBS containing 0.1% Tween-20 (PBST). Primary antibody incubation was performed overnight at 4 °C with rabbit anti-KCNV2 (1:1000) and mouse anti-GAPDH (1:1000) diluted in blocking buffer. After washing, membranes were incubated for 1 hour at room temperature with secondary antibodies: IRDye 800CW goat anti-rabbit (1:6000, LI-COR) and IRDye 680RD goat anti-mouse (1:10,000, LI-COR) and imaged using the ChemiDoc™ MP Imaging System (Bio-Rad Laboratories, Hercules, CA, USA). Band intensities were quantified using ImageJ software. KCNV2 signals were lane-wise normalized to GAPDH to account for potential variation in protein loading.

### Statistics

Statistical analysis and data visualization were performed using ggplot2[22], dplyr, readxl packages in R[23].

### Electroretinogram recordings

Full-field electroretinogram recordings were performed following in-house standards [24]. Specifically, recordings for scotopic and photopic conditions were conducted reflecting the standards of ISCEV [20] on ^E151X/E151X^ and WT mice of both sexes, at the age of 9-12 weeks. The mice were dark-adapted for at least 20 minutes before the experiment. The recordings were performed under shallow general anesthesia, which was induced using 5% isoflurane in 2 L/min O_2_ and maintained using 1.6-1.8 % of isoflurane in 0.8 L/min O_2_, respectively. Pupils were dilated by local application of a drop of Tropicamide 0.5% (Mydriaticum Stulln, manufacturer) and Phenylephrine 5% (Neosynephrin-POS 5%, manufacturer). Oxybuprocaine hydrochloride (4 mg/ml) was installed to the eye before placing of the electrodes. ERG recordings were performed using CELERIS ERG system (Diagnosys LLC, Lowell, MA, USA) which employs an integrated light guide stimulator and electrode placed on each eye and a ground electrode, attached to the tail. For rod response measurement light stimuli of flash energies ranging from 0.001-1 cd × s / m^2^ were presented to the dark-adapted mice whereas for cone response isolation light flashes of intensities ranging from 1-10 cd × s / m^2^ and frequencies of 10 Hz and 25 Hz were presented on a constant 30 cd/m^2^ background after >8 minutes of adaptation. Data acquisition was performed using the Espion software (Diagnosys) and digitized at 2kHz. ERG waveforms were analyzed to measure various parameters, such as the amplitude and latency of the a- and b-waves.

## 2. Results

### 2.1 Generation of a novel mousseline harboring the E151X mutation in the Kcnv2 gene

In order to develop a mouse model that would genetically more closely resemble the majority of cases of KCNV2-related retinopathy, we generated a mouse line carrying the early truncating E151X mutation (Figure 1A, c.583G>T), orthologous to the human E143X, in the Kcnv2 gene using CRISPR/Cas9 genome editing. The introduced mutation created a novel *BfaI* restriction site, enabling efficient genotyping of this mouse line by PCR amplification followed by *BfaI* digestion (Figure 1 B, C & D) Two founder animals were identified and back-crossed to C57Bl/6J mice for at least four generations before the line was used in experiments. Offspring from these matings were fertile and viable regardless of genotype or sex.

**Figure 1.**
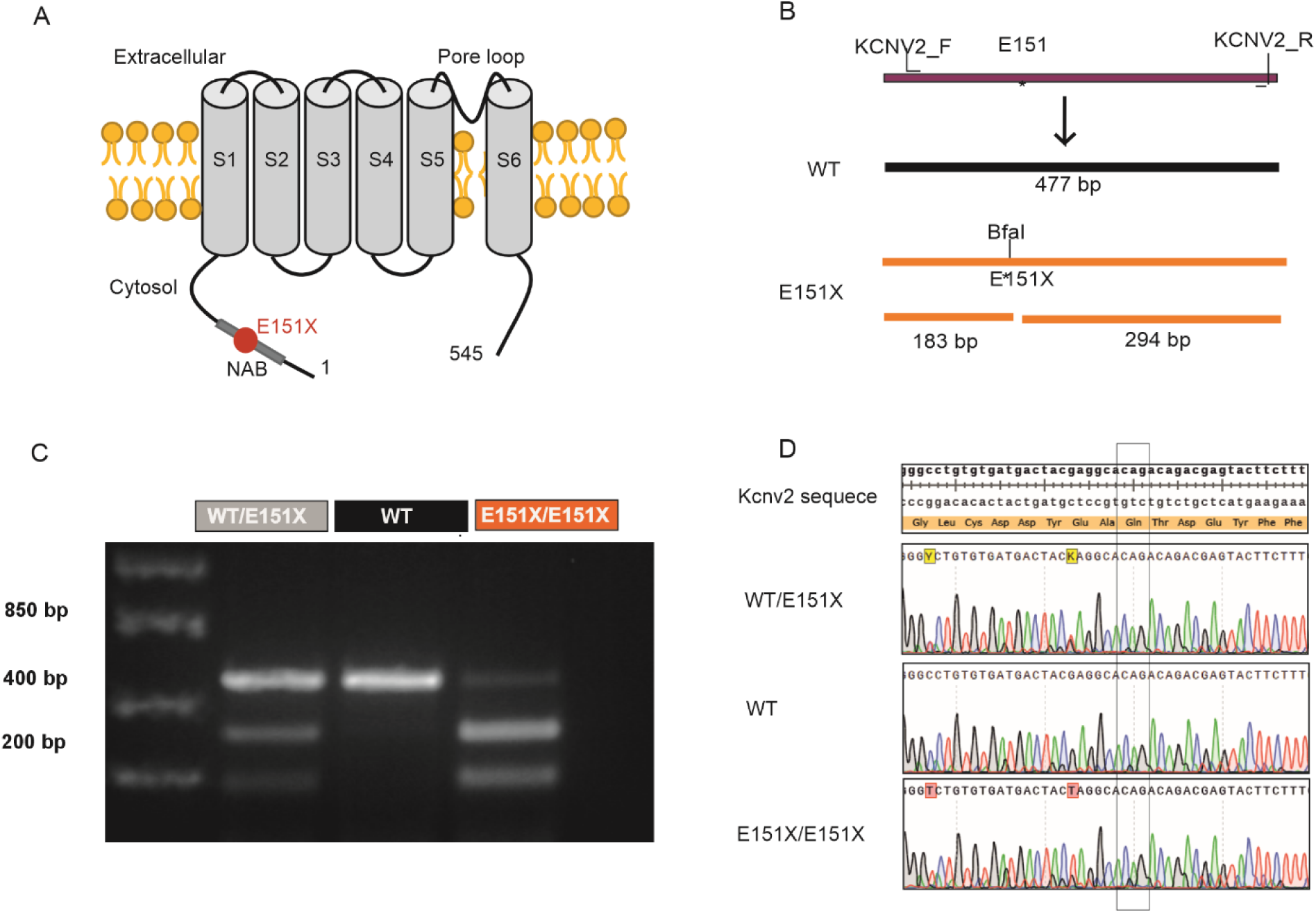
Generation and validation of the mutant mouse line using CRISPR/Cas9 and Sanger Sequencing. (A) Schematic representation of the mutation’s localization in the Kcnv2 tetramerization domain (NAB) (B) Genotyping strategy using PCR amplification followed by BfaI digestion. The restriction site is introduced only in the mutant allele, allowing selective cleavage. (C) Representative genotyping gel showing WT, heterozygous (Kcnv2^WT/E151X^) and homozygous mutant (Kcnv2^E151X / E151X^) animals and their corresponding band patterns. (D) Exemplary Sanger sequencing chromatograms of WT, Kcnv2 ^WT/E151X^, andKcnv2 ^E151X/E151X^ mice.

### 2.2 Kv8.2 is absent in the Kcnv2E151X mouse

Employing immunohistochemistry on retinal cryosections and western blot, we confirmed that the full-length protein was absent in these mice. Immunohistochemical staining of retinal cryosections resulted in distinct staining patterns in the Kcnv2^WT/WT^ tissues compared to the Kcnv2^E151X/E151X^ ones. In the WT samples, K_v_8.2 immunoreactivity was specifically localized to the photoreceptor inner segments, in full agreement with previous reports. In the Kcnv2^E151X/E151X^ retinal sections, we detected a complete absence of staining for K_v_8.2 in two mice and a weak residual K_v_8.2 signal for two other Kcnv2^E151X/E151X^ mice (Figure 2A). The residual signal can be possibly explained by the antibody recognizing the truncated K_v_8.2 protein, as the immunizing peptide used for antibody generation encompassed AA 1-163 of the K_v_8.2’s N-terminal cytoplasmic region, which is still present in the truncated protein. This faint signal was observed at the very tip of the inner segment of photoreceptors, but was markedly less intense compared to the WT tissues. To further evidence the absence of full-length K_v_8.2 in the Kcnv2E151X mouse model, we performed Western Blot analysis on retinal lysate samples from Kcnv2^E151X/E151X^ and WT littermates (n=3 each). Western blot analysis revealed the presence of full-length K_v_8.2 (65 kDa) in retinas from the WT mice. In contrast, no corresponding band was detected in the Kcnv2^E151X/E151X^ retinal lysates (Figure 2B). Taken together, these results confirm the effectiveness of the knockout strategy and validate the loss of full-length K_v_8.2 in our mouse line.

**Figure 2.**
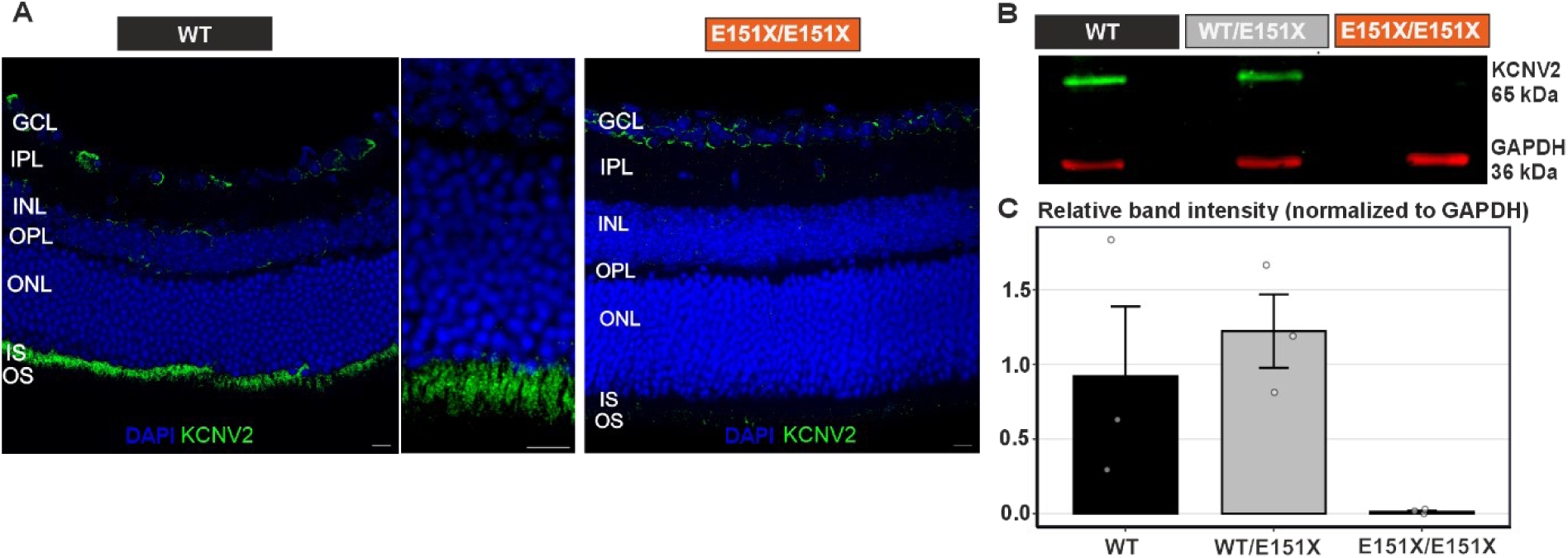
Expression of K_v_8.2 in WT, Kcnv2 ^WT/E151X^, and Kcnv2 ^E151X/E151X^ retinas. (A) Immunohistochemistry showing K_v_8.2 localization in retinal sections from WT (left panel), WT- zoomed (middle panel) and Kcnv2 ^E151X/E151X^ mice (right panel). K_v_8.2 is observed in the IS of the WT retina, while signal is absent in Kcnv2 ^E151X/E151X^ samples, confirming the loss of expression. (B) Representative Western blot of K_v_8.2 in retinal lysates from WT, Kcnv2 ^WT/E151X^, and Kcnv2 ^E151X/E151X^ littermates. GAPDH was used as a loading control. A 63 kDa band corresponding to K_v_8.2, was only detected in WT and heterozygous samples. (C) Quantification of K_v_8.2 band intensity normalized to GAPDH, showing comparable levels of protein expression in WT and Kcnv2 ^WT/E151X^ mice and complete absence of the protein in homozygous samples. Data are presented as mean ± SEM. n = 3 mice for each genotype.

### 2.3 Kcnv2E151X mice show a loss of cones and reduction in photoreceptors layers

To explore to what extent our new mouse line resembled the human disease phenotype, we first quantified cone loss at different ages by the means of quantifying cone arrestin (CArr) immunopositive somata in the outer nuclear layer. Kcnv2 ^E151X/E151X^ mice exhibited a continuous decline in CArr positive soma with age (slope = -0.18 CArr^+^ somata / week / Field of View [FoV], p = 0.013) while no such decline could be observed in WT littermates (slope = 0.00 CArr^+^ somata / week / FoV, p = 0.6, Figure 3). In addition to cone loss, a loss of photoreceptor cells in general in homozygous mutant mice was also observed, as evidenced by a decrease in rows of (DAPI^+^) nuclei in the ONL. Kcnv2 ^E151X/E151X^ mice lost -0.11 rows / week / FoV (p = 0.056) while the loss in WT littermates was substantially less pronounced (-0.06 rows / week / FoV, p = 0.185, Figure 3). We were not able to assess rod photoreceptor counts separately. It appears unlikely that the marked decrease in rows of ONL nuclei would be due to cone loss alone. We therefore conclude that also some rod loss is occurring in our Kcnv2 ^E151X^ mouse model.

**Figure 3.**
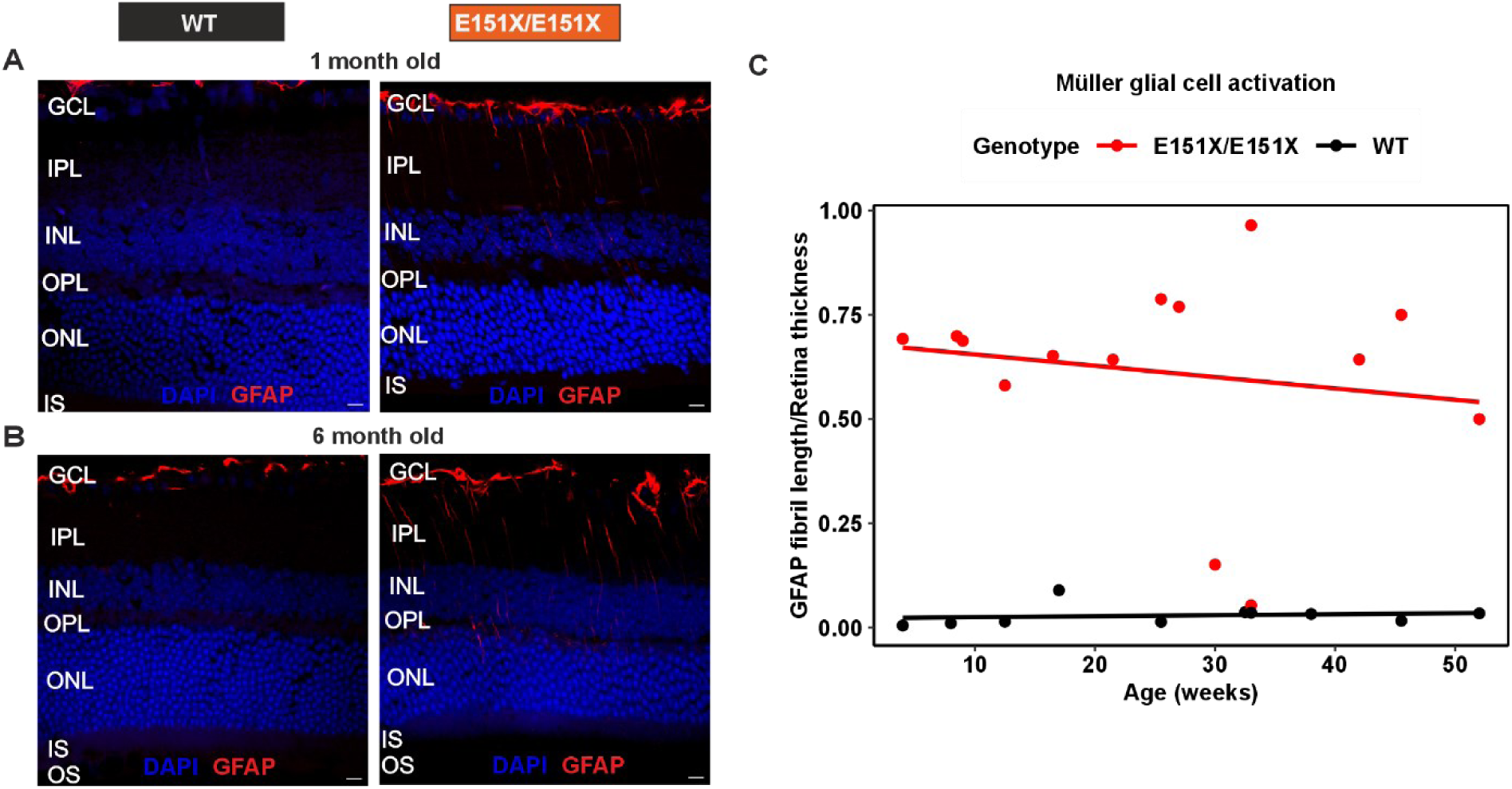
Müller glial activation in the Kcnv2E151X mouse model. (A) Representative retinal sections from 1-month-old WT and Kcnv2^E151X/E151X^ mice stained for glial fibrillary acidic protein (GFAP). At 1-month, in mutant mice GFAP immunoreactivity is already largely present as fibrils extending towards the inner retina, whereas stainings in age-matched WT mice no detectable glial activation. (B) GFAP staining in 6-month-old WT and Kcnv2 ^E151X/E151X^ retinas. Kcnv2^E151X/E151X^ mice display robust GFAP-positive radial fibrils extending towards the ONL, whereas WT retinas maintain GFAP localization remains at the GCL. (C) Quantification of GFAP fibril length normalized to retinal thickness as a function of age (weeks) in WT (black dots) and Kcnv2 ^E151X/E151X^ (red dots) mice. Kcnv2^E151X/E151X^ mutants show a significant upregulation in GFAP beginning at 4 weeks and persisting through more than 50 weeks of age, indicative of an early and sustained Müller glial activation, whereas WT values remain at baseline values. Each point represents one retina.

**Figure 4.**
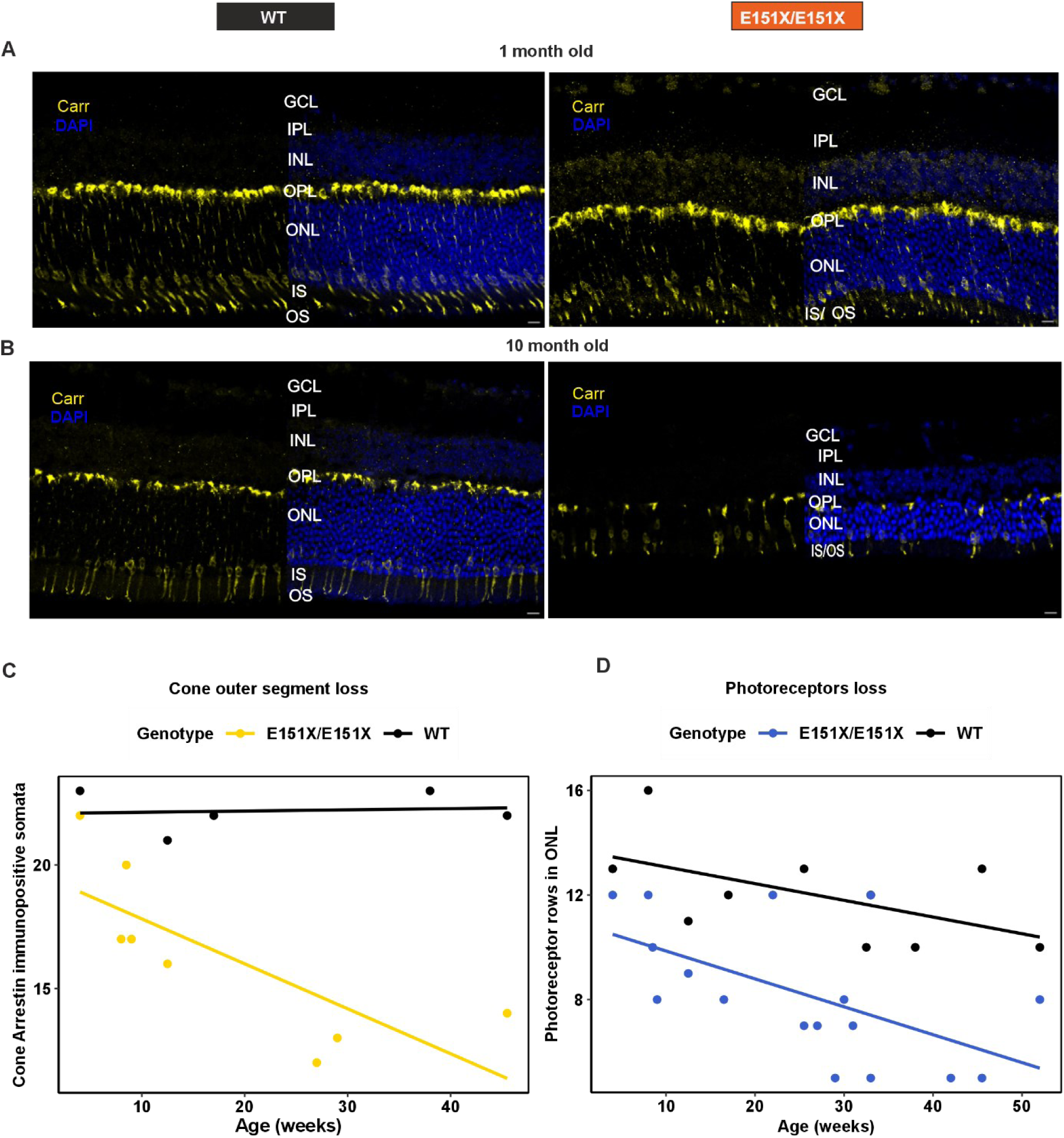
Progressive photoreceptor loss in Kcnv2^E151X/E151X^ mice. (A) Immunohistochemistry for cone arrestin in 1-month-old WT (left) and Kcnv2 ^E151X/E151X^ (right) retinas. Cone arrestin-positive cells are readily observed in both genotypes, with no marked deficit in the Kv8-mutant retinas. (B) Cone arrestin staining in 10-month-old WT and Kcnv2 ^E151X/E151X^ retinas. Kcnv2^E151X/E151X^ mice show a clear reduction in cone arrestin-positive cells compared to age-matched WT controls, indicating cone loss over time. (C) Quantification of cone arrestin-positive soma in the outer retinal layer across age. WT mice maintain stable cone arrestin expression, whereas Kcnv2^E151X/E151X^ mice exhibit a progressive decline, consistent with age-dependent cone loss. Each dot in the graph represents one mouse. (D) Outer nuclear layer (ONL) thickness measured across age shows progressive ONL thinning in Kcnv2^E151X/E151X^ mice, reflecting overall photoreceptor degeneration. WT mice show minimal ONL thinning over the same period.

### 2.4 GFAP is highly expressed in Kcnv2E151X mice

Reactive gliosis is commonly observed in retinal degenerations [25] and goes along with remodeling of the internal limiting membrane [26, 27] that alters the ability of certain Adeno- associated viruses (AAV) to penetrate the retina. We therefore used Glial Fibrillary Acidic Protein (GFAP) immunoreactivity as a marker for Müller glial activation. In Kcnv2 *^E151X/E151X^* mice, Müller glial cell activation was assessed by measuring the ratio of GFAP fibril length to retina thickness (Figure 2). The GFAP fibril length to retina thickness ratio was consistently higher in the mutant mice compared to the WT ones across various ages, with a noticeable difference starting from as early as 4 weeks old. Fibril length was mostly stable over ages, showed a minimal downward trend in Kcnv2^E151X/E151X^ mice (slope=0.003, p < 0.001). In contrast, no relevant activation of Müller glial cells could be observed in WT mice.

### 2.5 Visual function largely resembles Kcnv2 – associated retinopathy patients’ responses

To evaluate the electrophysiological characteristics of our mouse line, we performed full-field electroretinogram recordings in six Kcnv2^E151X/E151X^, seven WT and six Kcnv2^WT/E151X^ aged between 9-12 weeks. The rods response was assessed by measuring the amplitudes of a-wave and b-wave components under dark-adapted conditions at 0.001, 0.01, 0.1 and 1 cd × s/m^2^ stimuli intensities. The a-wave amplitudes were significantly smaller in Kcnv2^E151X/E151X^ mice compared to WT, when stimulated by 0.1 and 1 cd × s/m^2^ flashes. The heterozygous mice showed a phenotype that mostly resembled that of the WT mice, though, there was a slight, non-significant, reduction in amplitudes (Figure 5B). When contemplating the b-waves, the amplitudes in mutant mice were substantially and significantly lower than for WT mice upon dim flashes, while from 0.01 cd × s/m^2^ on, these increased to levels indifferent and sometimes exceeding those observed in WT mice (Figure 5C). We further analyzed the timing of the rod -pathway responses by examining the time to peak (implicit time) of the a- and positive b-wave under scotopic conditions. The Kcnv2^E151X/E151X^ mice displayed a delayed a-wave implicit time compared to WT mice, indicating a slower initial response of the rod photoreceptors to the light flashes (Figure 5D). Moreover, the a-wave troughs were broader in Kv8.2 ^E151X/E151X^ mice, as opposed to the WT and heterozygous ones (Figure 5A) Additionally, the b-wave implicit time was significantly prolonged in Kcnv2^E151X/E151X^ mice relative to WT mice, reflecting delayed post-phototransduction signaling in the rod pathway across an increasing light stimulus energy (Figure 5E). To evaluate cone function, we measured the photopic ERG responses under light-adapted conditions (Figure 6). The cone-mediated a-wave amplitudes were reduced in Kcnv2 ^E151X/E151X^ mice compared to WT mice. The responses were significantly smaller at 3 and 10 cd × s/m^2^ when compared to the WT group and only at 3 cd × s/m^2^ when compared to the heterozygous group (Figure 6 A & B). The decrease in amplitudes was also observed for the b-waves, which were markedly lower in the Kcnv2^E151X/E151X^ group from 3 cd × s/m^2^ onwards, compared to both- WT and heterozygous littermates. Analysis of oscillatory potentials revealed significant differences in OP amplitudes of Kcnv2^E151X/E151X^ mice. (Figure 7) Quantification of OP1, OP2, and OP3 at 0.1 cd × s/m² and analysis by multivariate linear regression a large reduction in OP amplitude in homozygous Kcnv2^E151X/E151X^, compared to both WT and heterozygous mice (Figure 7B).

**Figure 5.**
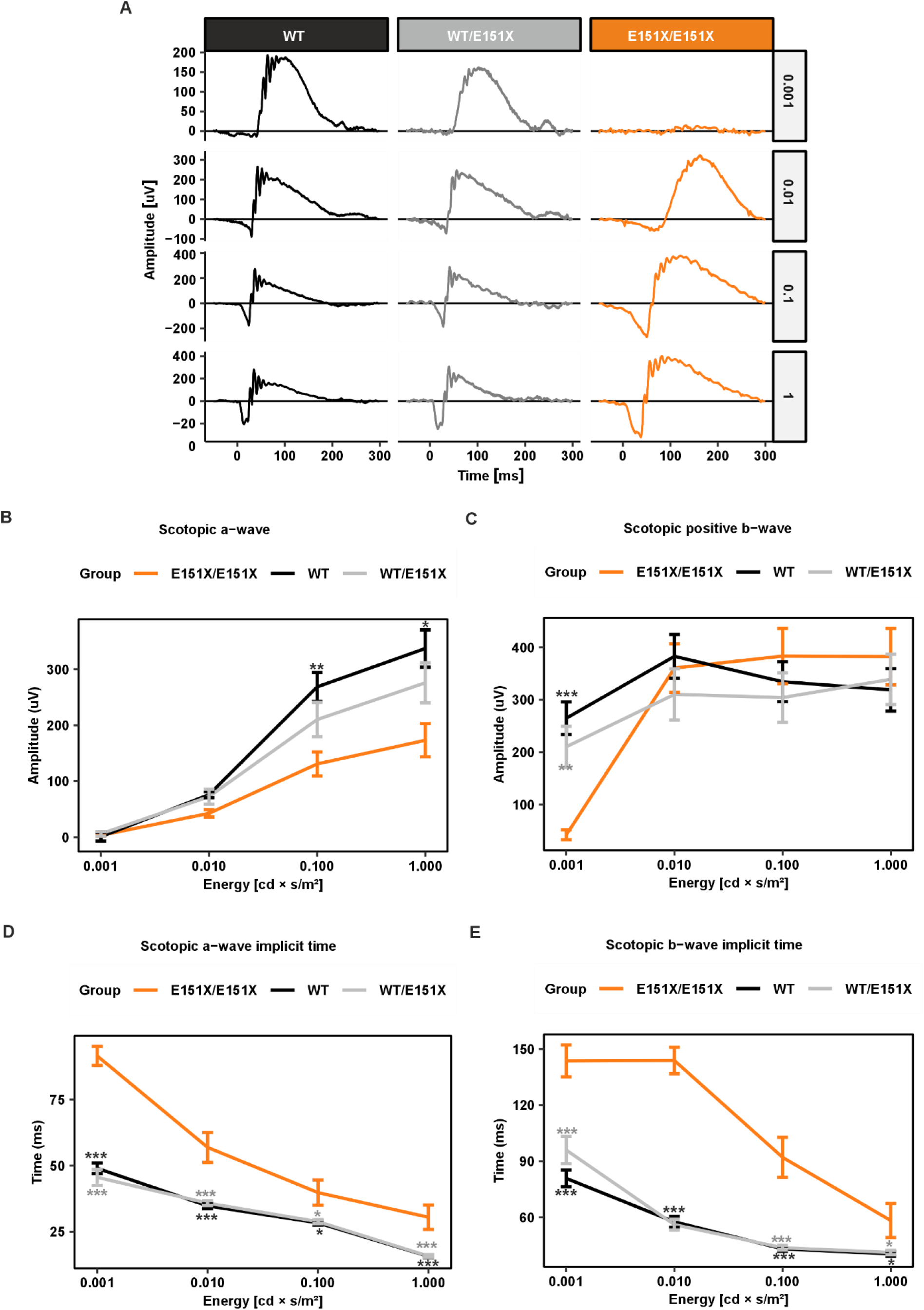
Scotopic ERG responses in WT, Kcnv2 ^WT/E151X^, and Kcnv2^E151X/E151X^ mice. (A) Representative ERG traces for each genotype in response to increasing flash energies (0.001, 0.01, 0.1, and 1 cd × s/m²), illustrating stimulus-dependent response characteristics. (B) Amplitude of the scotopic a-wave recorded from wild-type (WT), heterozygous (Kcnv2^WT/E151X^), and homozygous (Kcnv2^E151X/E151X^) mice. (C) Amplitude of the scotopic b-wave. (D-E) Implicit time of the scotopic b-wave and a-wave, showing timing differences in response kinetics across genotypes. Data for A-D are presented as mean ± SEM. Statistical significance was assessed using one-way ANOVA with post hoc Tukey test. n = 6-7 eyes per genotype. *p < 0.05, **p < 0.01, ***p < 0.001.

**Figure 6.**
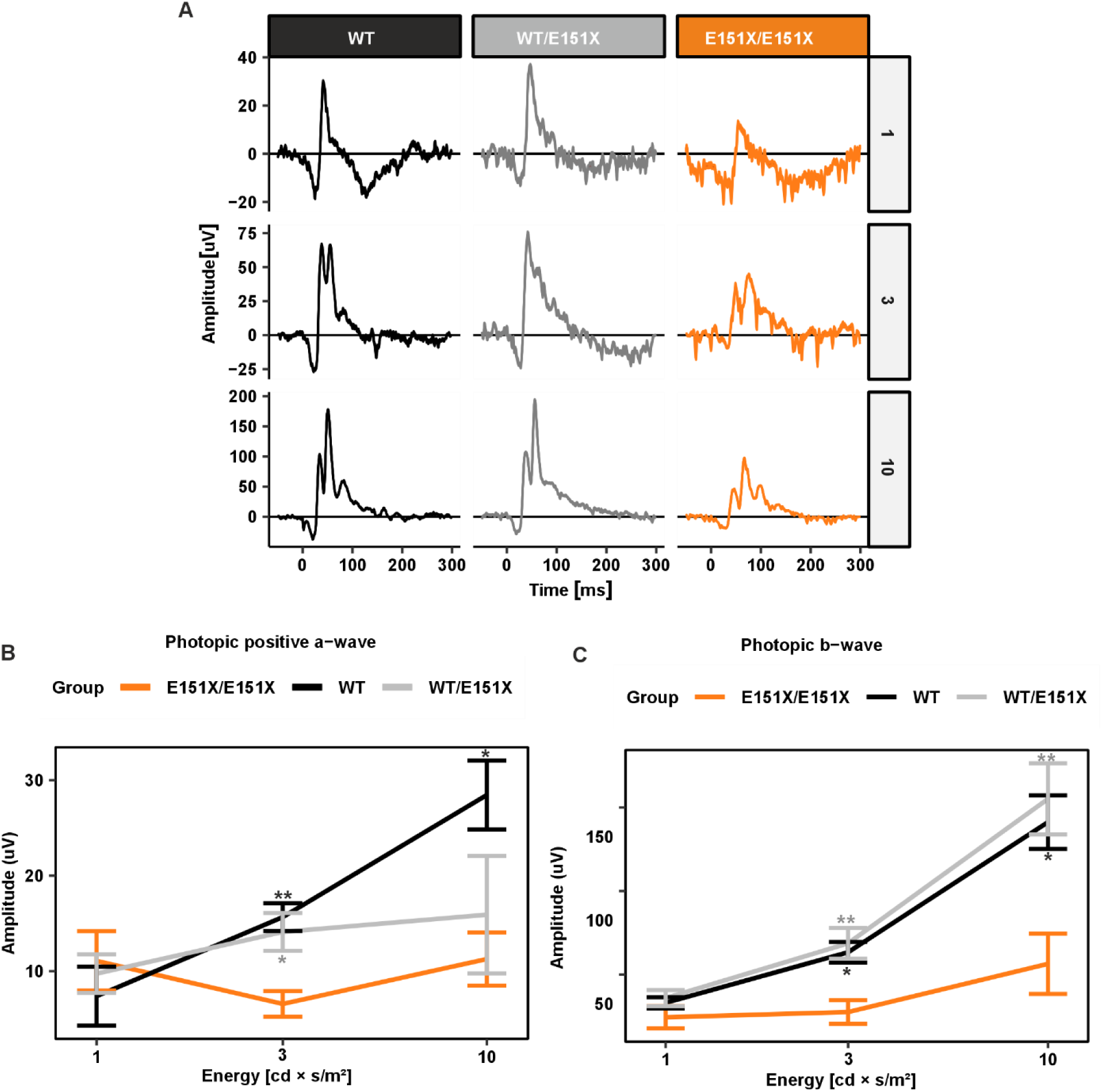
Photopic ERG responses in WT, Kcnv2 ^WT/E151X^, and Kcnv2 ^E151X/E151X^ mice. (A) Representative photopic ERG traces for each genotype in response to increasing flash energies (1, 3, and 10 cd × s/m²).(B) Amplitude of the photopic a-wave recorded from WT, Kcnv2 ^WT/E151X^, and Kcnv2^E151X/E151X^ mice under light-adapted conditions, reflecting cone photoreceptor activity. (C) Amplitude of the photopic b-wave for the same genotypes, indicating bipolar cell function integrity in the cone pathway. Data for B-C are shown as mean ± SEM. Statistical comparisons were performed using one-way ANOVA with Tukey’s post hoc. n = 5-7 eyes per group; *p < 0.05, **p < 0.01, ***p < 0.001.

**Figure 7.**
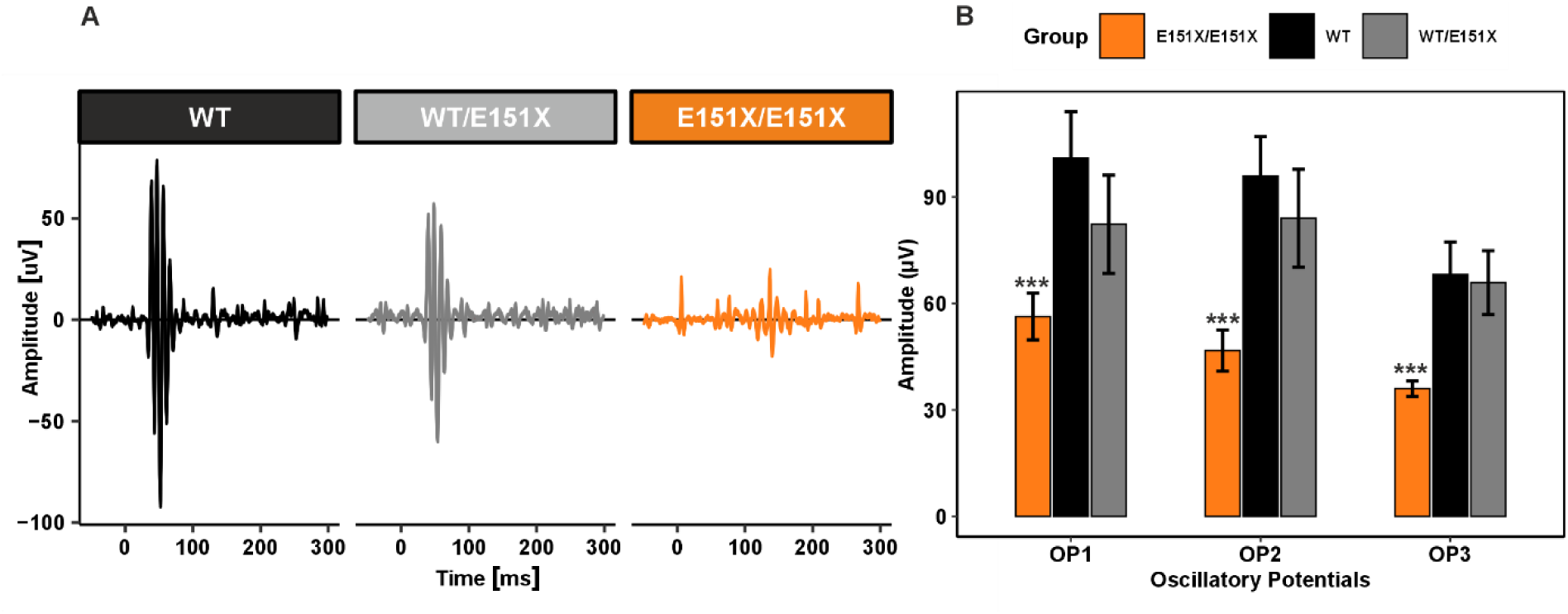
Representative oscillatory potential (OP) traces and quantification of OP amplitudes in WT, Kcnv2 ^WT/E151X^, and Kcnv2 ^E151X/E151X^ mice. (A) Representative oscillatory waveforms recorded from WT, Kcnv2 ^WT/E151X^, and Kcnv2 ^E151X/E151X^ mice, illustrating the characteristic reduction in OP amplitude in the Kcnv2 ^E151X/E151X^ group. (B) Quantitative comparison of OP1, OP2, and OP3 amplitudes across the three genotypes at 0.1 cd × s/m². Data are presented as mean ± SEM; statistical comparisons were performed with two-way ANOVA following multivariate regression with WT and OP1 as reference levels. Reduction of OP amplitudes were significant in homozygous mice, compared to both wild-type and heterozygous mice across all three OPs assessed. n = 5-7 eyes per group.

### 2.6 Interventional trials are feasible using similar functional outcome measure in mice and man

Having observed profound similarities between the human disease and the phenotype of the Kcnv2^E151X^ mouse, we reasoned that this mouse model might well be suited as a starting point for assessing efficacy of a hypothetical new treatment approach. We reasoned that given the structural stability of the rod system in particular and the characteristic electrophysiological changes, dark-adapted ERG metrics would be best suited for proof of concept. As nyctalopia is a common symptom in KCNV2-associated retinal dystrophy patients [15], it is moreover comparably straight forward to argue the clinical relevance of an improvement in dark-adapted ERG metrics towards a regulator. In this regard, having a single (or analog) metric that could be used in mouse and man might additionally facilitate translation.

The b-wave implicit time at dim stimulus intensities (DA 0.01 cd × s/m^2^) shows a strong contrast between wildtype and homozygous mice in our study (Figure 5 E) as well as between affected human individuals and controls[15]. We used this metric to simulate a power calculation for a hypothetical therapeutic trial for different treatment effect sizes and cohort sizes. We found that even for a moderate expected effect of only a 30% shift of b-wave implicit times towards normal values, a statistical power of 80% is reached at a sample size of 7 in mice and 12 in man (n per group, alpha = 0.05; Figure 8).

**Figure 8.**
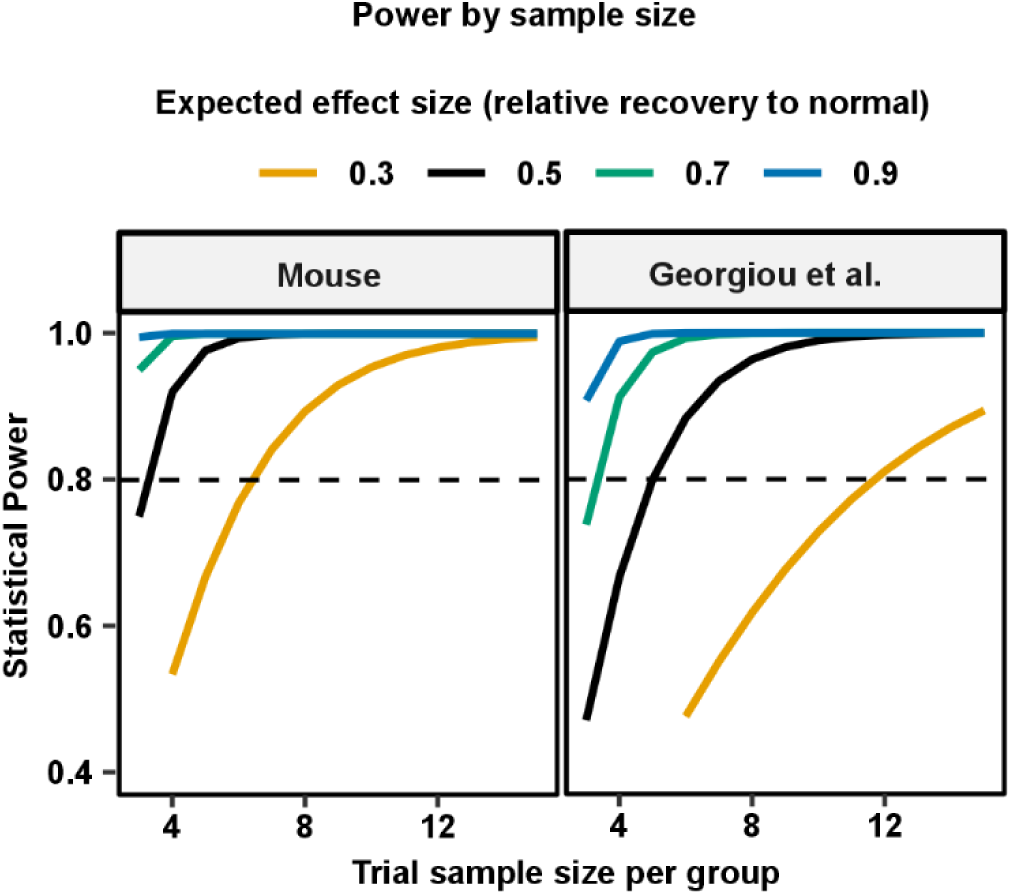
Statistical power analysis for a hypothetical therapeutic trial. Endpoint would be a 30%, 50%, 70% and 90% shift of DA 0.01 implicit b-wave times towards normal values, respectively. Statistical power (y-axis) is shown as a function of sample size (x-axis) using the dark-adapted (DA) 0.01 b-wave latency as the outcome measure. Power simulations based on data from our Kcnv2 mouse line (left panel) are compared to analysis on patient data reported by Georgiou et al. [15] (right panel).

## Discussion

In the present work we report on the generation and characterization of the Kcnv2^E151X^ mouse, carrying the orthologue of the common human E143X mutation. We found that the retinal phenotype of Kcnv2^E151X^ mirrors key characteristics of KCNV2-related retinopathy. Specifically, the mice lack expression of full-length K_V_8.2, undergo progressive cone loss and display the characteristic ERG pattern with delayed scotopic responses, disproportionate b-wave growth, and severely reduced photopic signals. These concordances indicate that the model accurately reflects human disease biology.

Two other mouse models for KCNV2-related retinopathy have been previously reported. [28, 29] As compared to those models, the mutation induced in the model described herein induces an early truncation in the middle the NAB domain as opposed to the (virtually) complete knock-out obtained in the previous models. [28, 29] E143X is the most common single variant associated with KCNV2-related retinopathy, accounting for 12.6% of all mutations linked to the disease [15]. Thus, on a genetical level, our model closely resembles this common disease-causing variant. Mutations within the NAB, which is relevant for efficient heteromerization of compatible K_V_ subunits [30] are overall common including several early truncation variants and 35.7% of all missense variants associated with the disease [15]. Thus, the model presented herein may be closely representative for a large proportion of affected individuals.

Smith et al had previously shown in cell line experiments that mechanistically distinct types of mutations may occur. Mutations in proximity to the NAB domain lead to production of homomeric K_V_2 channels that show currents that are larger and biophysically distinct when compared to the physiological K_V_8.2-K_V_2 heteromers (specifically: left-shifted voltage activation, faster inactivation). Pore mutations lead to the production of non-functional K_V_8.2-K_V_2 heteromers and a small portion of K_V_2 homomers, resulting in, most notably, a profound reduction in current amplitudes [16]. Although that study had only explored two mutations *neighboring* the NAB their data suggest that early mutations, like E151X (equalling the human E143X) might be susceptible to gene replacement therapy approaches or pharmacological strategies that primarily reduce current K_V_2-mediated current amplitudes. Notably, experiments on the functional consequences of the E151X mutation have yet not been published. It is also unclear what the consequence of the expression of aberrant Kcnv2- E151X protein is. Some Kcnv2- E151X mRNA might undergo nonsense-mediated decay [29]. Yet, to some extent Kcnv2- E151X protein will be produced, containing partial NAB domains that are compatible among Kcnv2- E151X and towards K_V_2 potentially resulting in intracellular accumulation or even sequestering K_V_2 partners. The fact that we detected some, mislocalized Kcnv2-immunoreactivity in some Kcnv2^E151X/E151X^ mice when using antibodies directed against the N-terminus is suggestive for a low degree of Kcnv2-E143X protein production – at least in some mice. How this influences the disease process could be studied in the mouse line presented herein, possibly by structural comparison to the available full knock-out lines.

Today there is robust evidence that K_V_8.2-K_V_2 heterotetramers substantially contribute to I_K,X_, including by electrophysiology and co-immunoprecipitation from native tissue [6]. Due to its similarity to the neuronal m-current the original hypothesis however had been that K_V_7 channels might contribute to I_K,X_ [8] and, in fact, K_V_7 channel blockers have shown to reduce I_K,X_ by ∼30 % (without those results reaching significance [6]. As K_V_7 channels are a proven pharmacological target, e.g. in the context of cardiovascular disease [31], their possible contribution to I_K,X_ still needs further evaluation and any possible contribution to I_K,X_ could be tested comparing the effects of K_V_7 modulators in the Kcnv2^E151X^ vs. wild-type mice.

The Kcnv2^E151X^ mouse phenocopies key characteristics of the human disease [15]. Specifically including dark-adapted b-waves that start subnormal at low, and showing accelerated progression towards higher stimulus energies. Also, the broadened a-wave trough and delayed b-wave peaks are present in the Kcnv2^E151X^ mouse. The supernormal b-wave at high stimulus intensities, that has initially been namesake for the disease, can only be observed occasionally in the Kcnv2^E151X^ mouse and on average remains on the upper margin of normality – just as in man [2, 15]. Notably, in this mouse model we observed a pronounced reduction photoreceptor layer thickness over time, which cannot be explained by a loss of cones alone, as these account only for 2.8 % of the total number of photoreceptors in the mouse retina. [32] Instead, we assume this represents concomitant rod loss. This is at least in partial in contrast to human disease, where the rod-dominated peripheral retina remains widely intact even in advanced disease stages [15, 33]. Our observation that rods degenerate over the disease course in mice is consistent with the observations made in the other two available mouse models [28, 29], suggesting that this is truly a difference between the human and the murine disease phenotype (as opposed to a mouse model artefact). Inamdar et al. highlight that in human the cone-exclusive fovea is surround by a ring of high rod density and hence structural macular affection – the hyperautofluorescent ring or even atrophy – as sometimes seen in Kcnv2-related retinopathy patients would necessarily involve rod rich areas[29]. Yet, it needs to be taken into consideration that that the highest rod densities are found clearly more eccentric than the structural alteration seen in KCNV2-related retinopathy [15, 34, 35]. We rather see parallels between the cone-to-Muller cell ratio and the location of the hyperautofluorescent ring[36]. In any case, rod loss in the mouse model exceeds that observed in human. Potential reasons may include distinct clearance of extracellular potassium by Muller cells, which might be least effective in the human (para-)fovea [28]. Or a specific susceptibility of cones in humans: Contrary to mice, human cones express a second K_V_2 Isoform, K_V_2.2, which is absent in rods. Thus, in humans I_K,x_ has a slightly distinct structural basis in rods and cones and the involvement of K_V_2.2 might lead to more unfavorable electrophysiological changes in these cells in the absence of Kcnv2. An alternative hypothesis to consider is that nocturnal mice utilize their rod system more than man, and this might, by some mechanism, lead to a more severe affection of the rods.

Nevertheless, the course of photoreceptor degeneration in the Kcnv2^E151X^ mouse is slow, with cones remaining immunohistochemically detectable until at least one year of age (Figure 4). This course temporally mirrors the clinical course in human patients[15, 37], indicating that there is ample room for therapeutic intervention. Under the assumption that normalization of ERG metrics would be good measures for therapeutic success we simulated power calculations for a hypothetical therapeutic trial. Indeed, we found that utilizing DA 0.01 b-wave latencies as outcome measure, statistical power could be reached with a feasible number of participants (Figure 8).

One interesting observation is that the physiological phenotype of the retina of heterozygous mice closely, but not entirely mirrors that of the WT mice, small differences in amplitudes, mostly not significant in a statistical context. This observation matches the clinical conclusions that the KCNV2-related retinopathy is a recessive disease. Yet, they also leave room for speculation on a minor haploinsufficiency phenotype. In this regard, refined electrophysiological protocols might be required to further characterize any such changes.

A particular strength of our report is that we provide longitudinal data for cone and global photoreceptor counts over the first year of life, a time span which would roughly correlate to the 40 years of human life. This information can prove helpful when aiming to model specific disease stages or planning treatment studies. Over the same timespan we also observe Muller glia activation. We observe that modest glia activation occurs early in the disease, but then remains roughly stable over the entire disease course. The latter indicates that no major retinal remodeling or glial seal formation seems to occur. Additionally, while sharing basic similarities with the other Kcnv2 mouse models,[28, 29] the model presented herein exhibits some features which parallel the human disease phenotype somewhat closer than those. Particularly, square-shaped/broaden a-waves are constantly observed in human upon bright DA stimuli and these are also present in our mouse line while this is not the case in others [28]. Similarly, cone loss is more pronounced in this model, than in the others[28, 29].

In summary, we have successfully created and characterized a novel mouse line for the KCNV2 -related retinopathy. This new model very closely resembles the morphological and electrophysiological changes described in human patients. Even though the mouse lacks a macula, the strongly conserved photoreceptor physiology and the matching diagnostic ERG patterns make this model a fit platform for testing treatment options and gaining deeper insight into the pathophysiology of the disease. A power calculation we provide on the basis of our model’s as well as published clinical data additionally demonstrates that such studies are feasible with a limited number of patients.

## Acknowledgments

The authors thank Burkhard Schütz and the Department of Anatomy for providing access to their cryotome.

## Declarations

### Author Contributions

*Participated in research design: NX, BD, ML*

*Conducted experiments: NX, SB, BD, ML*

*Performed data analysis: NX, BD*

*Wrote or contributed to the writing of the manuscript: NX, SB, BD, ML*

### Data Availability

The datasets generated during and/or analysed during the current study are available from the corresponding author on reasonable request.

## Funding

This work was supported by the German Research Foundation (LI 2846/5-1 and LI 2846/6-1 to ML).

### Ethics approval

This study did not involve human participants. Animal work was performed with approval of the relevant authorities and in accordance with the institutional Ethics Guidelines of Animal Care. Further details are provided in the Methods section.

### Consent to publish

Not applicable.

## Supplemental figures

**Figure S1.**
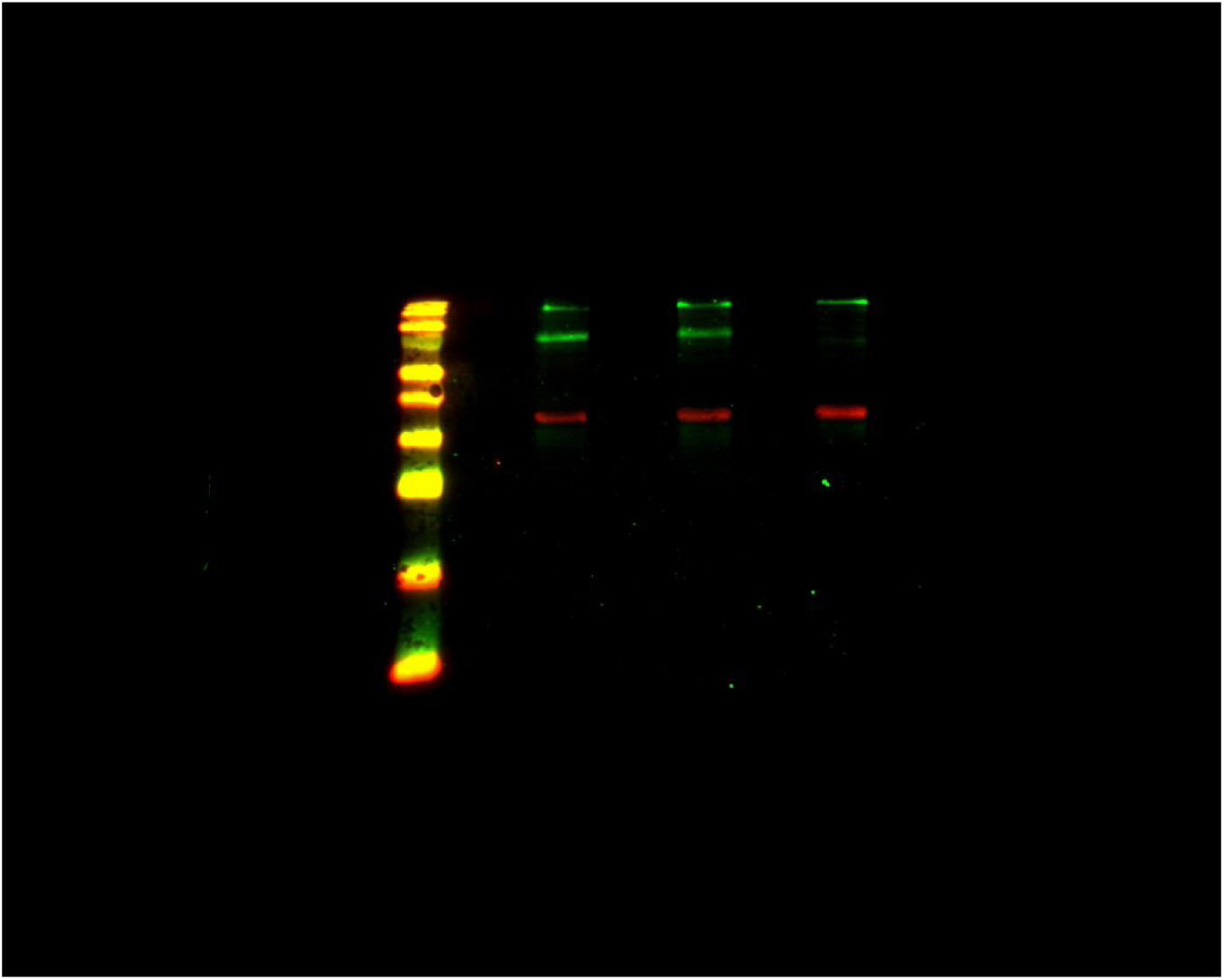
Raw Western blot for Kcnv2 in retinal lysates from three genotypes. Representative unprocessed Western Blot image showing Kcnv2 protein (green) and GAPDH (red) expression in retinal lysates from WT, heterozygous and E151X mice.

## References

1. Gouras, P., H.M. Eggers, and C.J. MacKay, Cone dystrophy, nyctalopia, and supernormal rod responses. A new retinal degeneration. Arch Ophthalmol, 1983. 101(5): p. 718–24.

2. Wu, H., et al., Mutations in the gene KCNV2 encoding a voltage-gated potassium channel subunit cause "cone dystrophy with supernormal rod electroretinogram" in humans. Am J Hum Genet, 2006. 79(3): p. 574–9.

3. Wissinger, B., et al., Cone dystrophy with supernormal rod response is strictly associated with mutations in KCNV2. Invest Ophthalmol Vis Sci, 2008. 49(2): p. 751–7.

4. Gill, J.S., et al., Progressive cone and cone-rod dystrophies: clinical features, molecular genetics and prospects for therapy. Br J Ophthalmol, 2019. 103(5): p. 711–20.

5. Sakti, D.H., et al., Natural history and biomarkers of KCNV2-associated retinopathy. Clin Exp Ophthalmol, 2024. 52(5): p. 528–544.

6. Gayet-Primo, J., et al., Heteromeric K(V)2/K(V)8.2 Channels Mediate Delayed Rectifier Potassium Currents in Primate Photoreceptors. J Neurosci, 2018. 38(14): p. 3414–3427.

7. Czirjak, G., Z.E. Toth, and P. Enyedi, Characterization of the heteromeric potassium channel formed by kv2.1 and the retinal subunit kv8.2 in Xenopus oocytes. J Neurophysiol, 2007. 98(3): p. 1213–22.

8. Barnes, D.J.B.a.S., Characterization of a Voltage-Gated K+ Channel That Accelerates the Rod Response to Dim Light. Neuron, 1989. **Volume** 3( Issue 5): p. Pages 573-581.

9. Khan, A.O., M. Alrashed, and F.S. Alkuraya, ’Cone dystrophy with supranormal rod response’ in children. Br J Ophthalmol, 2012. 96(3): p. 422–6.

10. Vincent, A., et al., Phenotypic characteristics including in vivo cone photoreceptor mosaic in KCNV2-related "cone dystrophy with supernormal rod electroretinogram". Invest Ophthalmol Vis Sci, 2013. 54(1): p. 898–908.

11. Robson, A.G., et al., "Cone dystrophy with supernormal rod electroretinogram": a comprehensive genotype/phenotype study including fundus autofluorescence and extensive electrophysiology. Retina, 2010. 30(1): p. 51–62.

12. Alexander, K.R. and G.A. Fishman, Supernormal scotopic ERG in cone dystrophy. Br J Ophthalmol, 1984. 68(2): p. 69–78.

13. de Guimaraes, T.A.C., et al., KCNV2-associated retinopathy: genotype-phenotype correlations - KCNV2 study group report 3. Br J Ophthalmol, 2024. 108(8): p. 1137–1144.

14. Ben Salah, S., et al., Novel KCNV2 mutations in cone dystrophy with supernormal rod electroretinogram. Am J Ophthalmol, 2008. 145(6): p. 1099–106.

15. Georgiou, M., et al., KCNV2-Associated Retinopathy: Genetics, Electrophysiology, and Clinical Course-KCNV2 Study Group Report 1. Am J Ophthalmol, 2021. 225: p. 95–107.

16. Smith, K.E., et al., Functional analysis of missense mutations in Kv8.2 causing cone dystrophy with supernormal rod electroretinogram. J Biol Chem, 2012. 287(52): p. 43972–83.

17. Sergouniotis, P.I., et al., High-resolution optical coherence tomography imaging in KCNV2 retinopathy. Br J Ophthalmol, 2012. 96(2): p. 213–7.

18. Alghadban, S., et al., Electroporation and genetic supply of Cas9 increase the generation efficiency of CRISPR/Cas9 knock-in alleles in C57BL/6J mouse zygotes. Sci Rep, 2020. 10(1): p. 17912.

19. Kinder, L. and M. Lindner, Expression of Osteopontin in M2 and M4 Intrinsically Photosensitive Retinal Ganglion Cells in the Mouse Retina. Invest Ophthalmol Vis Sci, 2025. 66(2): p. 14.

20. Carpentiero, E., et al., Interaction between native and prosthetic visual responses in optogenetic visual restoration. JCI Insight, 2025. 10(11).

21. Schindelin, J., et al., Fiji: an open-source platform for biological-image analysis. Nat Methods, 2012. 9(7): p. 676–82.

22. Wickham, H., ggplot2: Elegant Graphics for Data Analysis. 2016: Springer-Verlag.

23. R Core Team. R: A language and environment for statistical computing. 2021; Available from: https://www.R-project.org.

24. Lindner, M., The ERGtools2 package: A Toolset for Processing and Analysing Visual Electrophysiology Data. 2024.

25. Hippert, C., et al., Muller glia activation in response to inherited retinal degeneration is highly varied and disease-specific. PLoS One, 2015. 10(3): p. e0120415.

26. Ahuja, S., et al., rd1 mouse retina shows imbalance in cellular distribution and levels of TIMP-1/MMP-9, TIMP-2/MMP-2 and sulfated glycosaminoglycans. Ophthalmic Res, 2006. 38(3): p. 125–36.

27. Kolstad, K.D., et al., Changes in adeno-associated virus-mediated gene delivery in retinal degeneration. Hum Gene Ther, 2010. 21(5): p. 571–8.

28. Hart, N.S., et al., The Role of the Voltage-Gated Potassium Channel Proteins Kv8.2 and Kv2.1 in Vision and Retinal Disease: Insights from the Study of Mouse Gene Knock-Out Mutations. eNeuro, 2019. 6(1).

29. Inamdar, S.M., et al., Differential impact of Kv8.2 loss on rod and cone signaling and degeneration. Hum Mol Genet, 2022. 31(7): p. 1035–1050.

30. Yu, W., J. Xu, and M. Li, NAB domain is essential for the subunit assembly of both alpha-alpha and alpha-beta complexes of shaker-like potassium channels. Neuron, 1996. 16(2): p. 441–53.

31. Mackie, A.R. and K.L. Byron, Cardiovascular KCNQ (Kv7) potassium channels: physiological regulators and new targets for therapeutic intervention. Mol Pharmacol, 2008. 74(5): p. 1171–9.

32. Jeon, C.J., E. Strettoi, and R.H. Masland, The major cell populations of the mouse retina. J Neurosci, 1998. 18(21): p. 8936–46.

33. Zobor, D., et al., Rod and cone function in patients with KCNV2 retinopathy. PLoS One, 2012. 7(10): p. e46762.

34. Sato, T., et al., Clinical course of two siblings with potassium voltage-gated channel modifier subfamily V member 2 (KCNV2)-associated retinopathy. Doc Ophthalmol, 2024. 148(3): p. 173–182.

35. Curcio, C.A., et al., Human photoreceptor topography. J Comp Neurol, 1990. 292(4): p. 497–523.

36. Masri, R.A., et al., Immunohistochemistry and Spatial Density of Muller Cells in the Human Fovea. Invest Ophthalmol Vis Sci, 2025. 66(2): p. 46.

37. Georgiou, M., et al., KCNV2-Associated Retinopathy: Detailed Retinal Phenotype and Structural Endpoints-KCNV2 Study Group Report 2. Am J Ophthalmol, 2021. 230: p. 1–11.

